# BlueRecording: A pipeline for the efficient calculation of extracellular recordings in large-scale neural circuit models

**DOI:** 10.1101/2024.05.14.591849

**Authors:** Joseph Tharayil, Jorge Blanco Alonso, Silvia Farcito, Bryn Lloyd, Armando Romani, Elvis Boci, Antonino Cassara, Felix Schürmann, Esra Neufeld, Niels Kuster, Michael Reimann

**Affiliations:** Blue Brain Project, École polytechnique fédérale de Lausanne (EPFL) Campus Biotech, Geneva, Switzerland; Foundation for Research on Information Technologies in Society (IT’IS), Zurich, Switzerland

## Abstract

As the size and complexity of network simulations accessible to computational neuroscience grows, new avenues open for research into extracellularly recorded electric signals. Biophysically detailed simulations permit the identification of the biological origins of the different components of recorded signals, the evaluation of signal sensitivity to different anatomical, physiological, and geometric factors, and selection of recording parameters to maximize the signal information content. Simultaneously, virtual extracellular signals produced by these networks may become important metrics for neuro-simulation validation. To enable efficient calculation of extracellular signals from large neural network simulations, we have developed *BlueRecording*, a pipeline consisting of standalone Python code, along with extensions to the Neurodamus simulation control application, the CoreNEURON computation engine, and the SONATA data format, to permit online calculation of such signals. In particular, we implement a general form of the reciprocity theorem, which is capable of handling non-dipolar current sources, such as may be found in long axons and recordings close to the current source, as well as complex tissue anatomy, dielectric heterogeneity, and electrode geometries. To our knowledge, this is the first application of this generalized (i.e., non-dipolar) reciprocity-based approach to simulate EEG recordings. We use these tools to calculate extracellular signals from an *in silico* model of the rat somatosensory cortex and hippocampus and to study signal contribution differences between regions and cell types.

## 1 Introduction

The source of key signals measured in neuroscience is the transmembrane current generated by the electrical activity of neurons. Computational investigation of such signals using compartmental models requires: (1) a model and simulator that calculate the membrane currents, and (2) an estimation of how each compartment’s current affects the signal. For (2), typically simplifying approaches have been used. Simply comparing model output to reconstructed current dipoles relies on accurate and reliable source reconstruction from EEG data using inverse solution methods [1] – something that is difficult to achieve because of the poorly conditioned inverse problem and the complex impact of volume conduction in the heterogeneous head anatomy on EEG signal formation. Point-source and line-source approaches approximate the properties of the different tissues as homogeneous [2] [3] [4]. They are therefore only valid where electrodes are within the brain tissue, where some degree of homogeneity of the nearby dielectric environment can be assumed. The commonly employed simplified variant of the reciprocity theorem approximates individual neurons or groups of neurons as dipolar sources. It is therefore only valid when electrodes are sufficiently far away from said neurons [5].

Our goals in this work were to: provide a solution to (2) that is equally valid for both proximal and distal electrical measurements; provide flexibility to researchers with respect to the approaches to both (1) and (2) they want to employ; and to support very large electrophysiological models by enabling online and efficient computation.

We use the general form of the reciprocity theorem developed by Plonsey [6], which supports full compartmental resolution of the neuronal models and takes into account the dielectric properties of all tissues, to calculate the impact of each compartment on the signal. It can be employed for EEG, ECoG and LFP, and is capable of dealing with arbitrarily shaped electrode geometries. The impact of each compartment on the signal is stored in what we call a weights file. We then implemented extensions to the Neurodamus simulation control software [7] and the CoreNEURON simulation engine [8] to consume a weights file to calculate the signal during a simulation run and store the result for later analysis. Together, we call the applications to calculate weights files and the extensions to the simulator the BlueRecording framework.

The formalism of a standardized weights file ensures flexibility with respect to objective (2). A user can easily implement a different solution so long as it provides a valid weights file. Since largescale simulations are required for the EEG signal, the ability to use the highly efficient CoreNEURON simulation engine regardless of dielectric model is a significant advantage. In addition, BlueRecording is compatible with any circuit defined in the SONATA format [9], making it easily portable between different simulators, and therefore highly flexible with respect to objective (1) as well.

In this paper, we apply BlueRecording microcircuit models developed by the Blue Brain Project (BBP). The BBP somatosensory cortex model consists of ∼4.2 million reconstructed neurons with accurate morphologies, optimized physiological properties, and algorithmically generated connectivity [10]. The BBP hippocampus CA1 model consists of ∼456,000 mophologically detailed neurons, with properties and connectivity generated in a similar manner to [10], along with virtual innervation by Schafer collaterals and cholinergic modulation [11]. The support for spatially extended and morphologically highly detailed BBP microcircuits is a particular advantage when studying EEG signals. This is demonstrated here by comparing virtual LFP, EEG, and ECoG, and identifying various contributions to the signals.

## 2 Design and implementation

The extracellular signals are calculated online in the NEURON simulation environment. Due to the linear nature of Maxwell’s equation, the signal at a particular electrode (relative to another electrode or ground) is amenable to the form

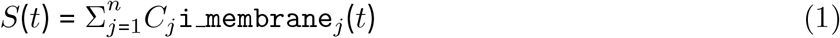

where `i_membrane` is the total compartmental transmembrane current (i.e., the sum of the ionic and capacitive currents, as described in [2]), *j* is the index of each neural compartment in the circuit, and *C* is a scaling factor (‘weight’), the calculation of which is discussed below. The equation takes a similar form as the commonly employed lead-field matrices, only that the latter apply to dipole source distributions instead of transmembrane currents. The simulator supports simultaneous recording from an arbitrary number of electrodes.

### 2.1 Calculation of coefficients

#### 2.1.1 Generalized reciprocity approach

Following Plonsey [12], it can be shown that *∰*_*U*_ *I*_*U*_ (*t*) ⋅ Φ_2_*dU* = *∯*_*S*_ Φ_1_ (*t*) ⋅ *KdS* where *I*_*U*_ (*t*) is a volume current source in an arbitrary dielectric environment *U* (in the case of a neural current source, *I*_*U*_ (*t*) is the set {`i_membrane`_*j*_ (*t*) ∀ *j*}), Φ_1_(*t*) is the potential resulting from to that current source, *K* is a surface current distribution on the dielectric material, and Φ_2_ is the potential resulting from this surface current distribution.

When *K* consists of a pair (*a,b*) of quasi-perfect electric conductors (e.g., metallic EEG electrodes) providing a (virtual) current *J*, the situation simplifies to *∰*_*U*_ *I*_*U*_(*t*) ⋅Φ_2_*dU* = *J* ⋅ (Φ_1,*a*_ −Φ_1,*b*_) = *J* ⋅ *V* (*t*), where *V* (*t*) = Φ_1,*a*_ 1(*t*) − Φ_1,*b*_ corresponds to the measured signal.

For a neurite with electrodes sufficiently far away, we can treat each nodal current `i_membrane`_*j*_ (*t*) ∈ *I*_*U*_ (*t*) as point current sources at points 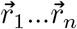, respectively. Thus, the left side of the equation reduces to 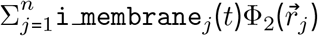 and the potential between the two electrodes in this situation becomes

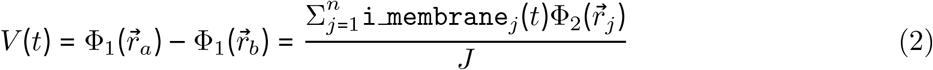

The coefficients *C*_*j*_ used to calculate the extracellular signal as described in Equation 1 are therefore given by

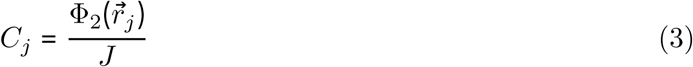

In order to calculate 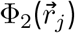, a finite element method (FEM) simulation of a head model with a recording electrode array is performed, where a (virtual) current is applied between a pair of recording electrodes, and the potential field is interpolated at the locations of the neural segments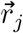. Information about the FEM rat head model setups created for this study is provided in Appendix A.

#### 2.1.2 Dipole-based reciprocity approach

If we take the Taylor series expansion of 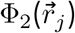 around 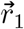, we obtain, to first order

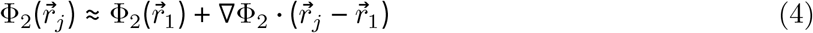

Substitution Equation 4 into Equation 2 and rearranging, we obtain

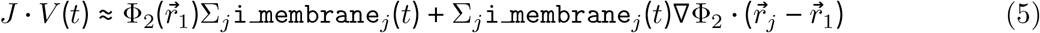

For a neural current source, Σ_*j*_`i_membrane`_*j*_ = 0 by Kirchhoff’s Law. Thus,

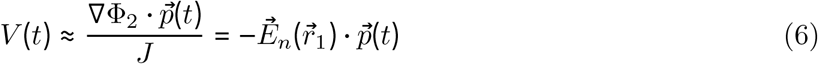

where 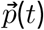 is the current dipole calculated per individual discretized neuron *N* at each time step as

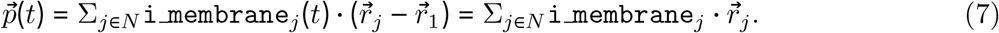

The normalized 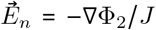 is the ‘lead-field’. 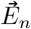 can be computed using a FEM model, as described in Section A, and is interpolated at the individual soma locations. Substituting into Equation 1 and rearranging, we find that

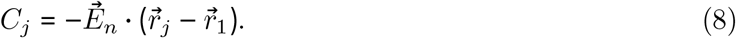

While the choice of 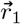 is not relevant from an analytical perspective, it can be numerically advantageous to place it near the neuron center.

#### 2.1.3 Calculation of coefficients with line-source approximation

Unlike EEG or ECoG electrodes, LFP electrodes are small and located within the cortical tissue. If the LFP electrode can be treated as point-like, the tissue within the relevant environment is homogeneous and isotropic, and the ground reference sufficiently distant to be treated as being at infinity and omnidirectional, the computation of an ‘exposing’ potential required by the reciprocity theorem can be avoided and instead the line-source approximation can be used to compute the coefficients for LFP calculations without need for FEM simulation.

According to the line-source approximation [13], the contribution to the electric potential at a point electrode location of a neural segment situated in a homogeneous, isotropic environment is given by 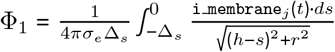, where Δ_*s*_ is the length of the segment, *h* is the distance from the near end of the segment to the recording electrode in the axial direction, and *r* is the absolute value of the distance from the near end of the electrode to the electrode in the perpendicular direction. Evaluating the integral analytically and substituting into Equation 1, we find that:

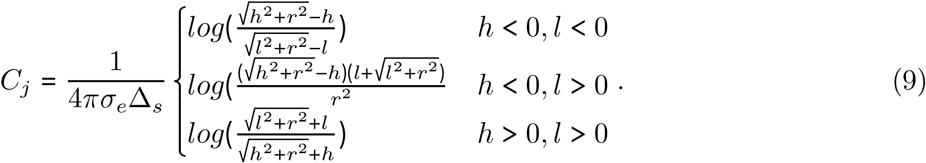

With (*x*_1_, *y*_1_, *z*_1_) and (*x*_2_, *y*_2_, *z*_2_) as the endpoints of the segment and (*x, y, z*) as the location of the electrode,

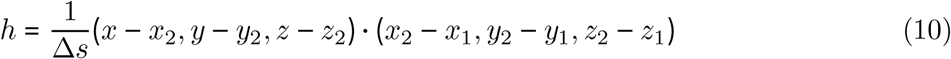

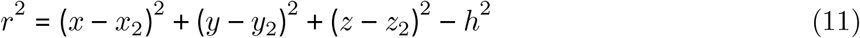

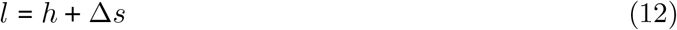

#### 2.1.4 Calculation of coefficients with point-source approximation

A further simplification can be made (in addition to the assumptions of an infinite homogeneous medium, a point-like electrode, and reference at infinity) by treating each neural compartment as a point rather than a line. In this case, the potential at the recording electrode is given by

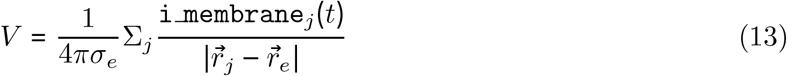

where 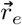 is the position of the electrode and 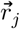 is the position of the compartment. Substituting into Equation 1 we obtain:

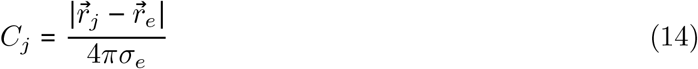

#### 2.1.5 Choice of voltage and spatial reference

The extracellular signals produced using the reciprocity approaches are gauge-independent with respect to the potential field (i.e., the location to which a potential of 0V is assigned is arbitrary). This is because the sum of membrane currents of the compartments over a neurons must be zero. Thus, any constant added to the compartment weights does not affect the signal, and only the difference in compartment weights matters. For better visualization, we can therefore add an offset to the compartment weights in each neuron such that the minimum weight per neuron is 0.

#### 2.1.6 Numerical considerations

Calculation of the extracellular signal involves summation over a large number of terms that typically compensate, potentially leading to important numerical errors as a large number of digits can be significant. Because of gauge freedom and current conservation, this can be mitigated by selecting good reference values for *r* and Φ_2_, and if that proves insufficient, by using a compensation algorithm such as Kahan summation. Our implementation does not use Kahan summation, as we have not found it to be necessary for the current study. However, awareness that compensated summation might be required in other situations, is necessary.

### 2.2 Workflow

The workflow for calculating coefficients and running an online extracellular recording simulation with BlueRecording is as follows (see as well Fig. 1). Tools used for each step are listed in parentheses:

**Figure 1:**
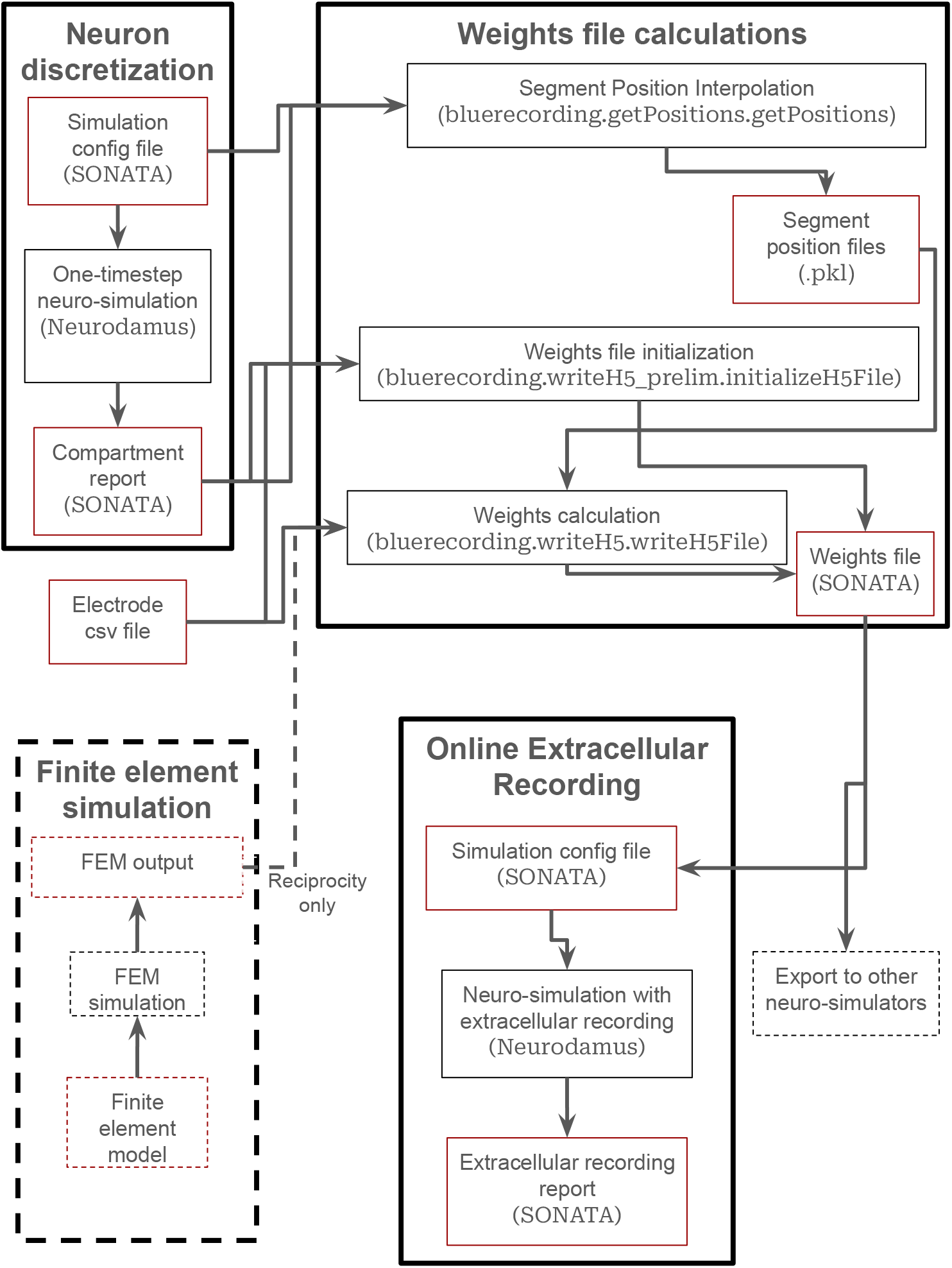
Workflow for the BlueRecording Pipeline. Input and output files are shown in red boxes, while processes are shown in black boxes. Dashed lines around the FEM simulation step indicate that it is only used for reciprocity-based calculations

1. **Calculate Segment Positions:** Segment positions are calculated in two steps
  a. **Run Neuro-Simulation (Neurodamus):** A minimal (at least one time step) neuro-simulation is run to produce a compartment report that reveals how the neurons are discretized.
  b. **Interpolate segment positions (**`bluerecording.getPositions.getPositions()`): The compartment report and information from the simulation configuration file are used to interpolate the segment positions from the 3d coordinates in the morphology file, which are not necessarily aligned to the discretized compartments
2. **Create electrode csv file (Manual)** See Section 2.3.2
3. **Initialize Electrode Weights file (**`bluerecording.writeH5 prelim.initializeH5File()`): This step reads metadata from the electrode csv file and the discretization from the compartment report, and writes both to the weights file. See Sections 2.3.1 for details on the file format and 2.3.3 for details on writing the file
4. **Calculate weights:**
  a. **Reciprocity-based methods**
    i. **FEM simulations (Any FEM solver):** A FEM simulation is run to calculate the potential field generated by running a current between a recording electrode and a reference electrode. The results (see Appendix C) are exported.
    ii. **Interpolation of weights (**`bluerecording.writeH5.writeH5File()`): From the exported results, the weights are calculated by interpolating the potential field at the neural compartment locations (for the generalized reciprocity approach) or by taking the scalar product of the E field at the center of the neuron and the segment position vectors (for the dipole-reciprocity approach).
  b. **Analytic calculations (**`bluerecording.writeH5.writeH5File()`): For the line-source and point-source approximations, the weights are calculated analytically, without requiring FEM simulations.
5. **Run neuro-simulations (Neurodamus or alternative neuro-simulator)** While it is intended that simulations be run using Neurodamus as described in Section 2.3.4, other simulators may be extended to also support the weights file created in BlueRecording.

### 2.3 Implementation details

#### 2.3.1 Extension to SONATA-format

We defined an extension to the SONATA format [9] to store the coefficients *C*_*j*_. Briefly, the SONATA format treats each neuron in the network as a **node**, and nodes are organized into **populations**. The coefficients are stored in an H5 file (Table 1), which also includes metadata on the type and location (both in Cartesian space and with respect to the anatomy) for each electrode. For each population, the coefficients for all nodes and all electrodes are listed in a single array; for each node, the coefficients (one per compartment) are listed consecutively. For each population, there is also a dataset listing the IDs of each node therein, as well as the index in the coefficient array at which the coefficients for that node begin.

**Table 1:**
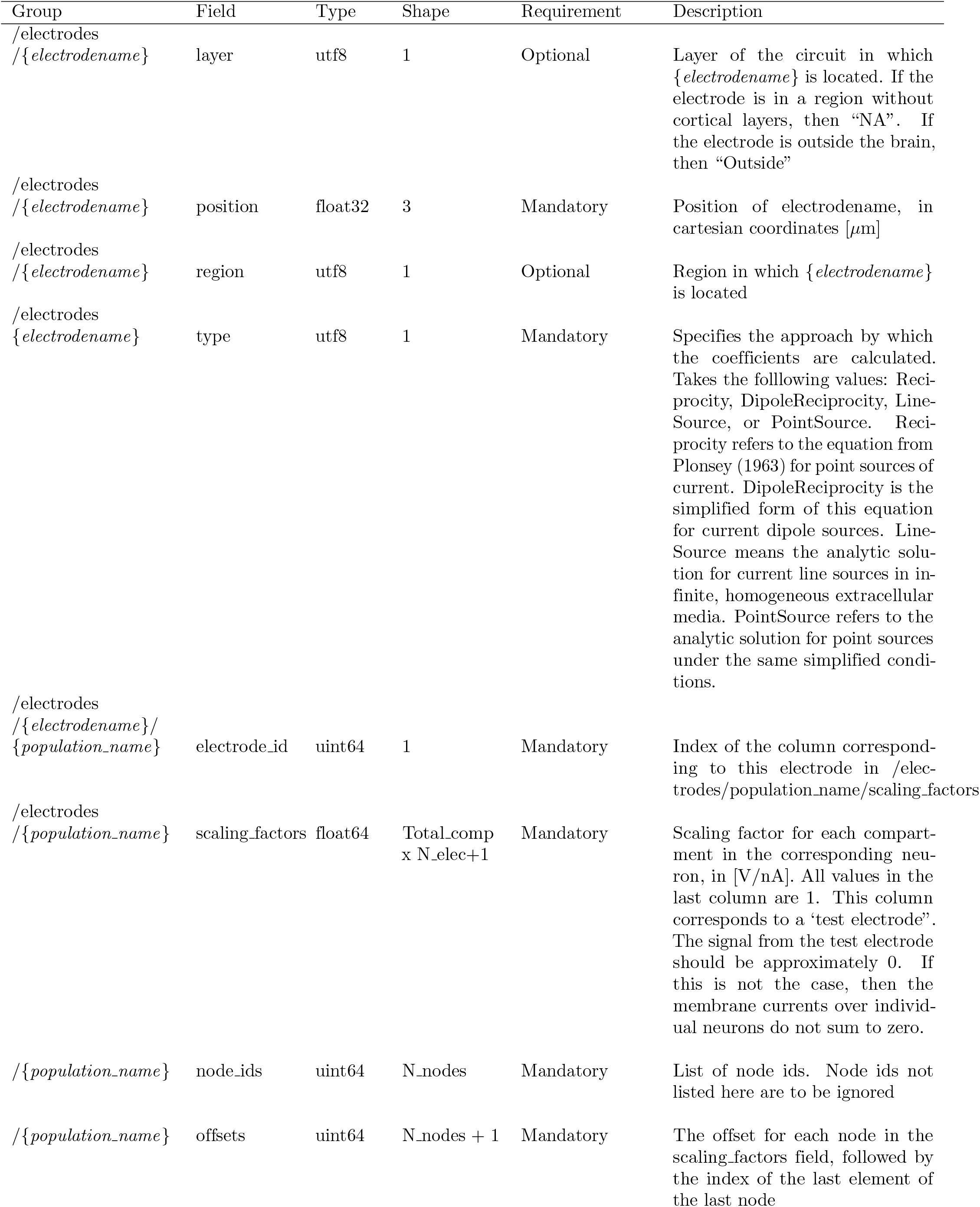
The format of the weights file.

#### 2.3.2 Electrode array file format

The electrode array for which the coefficients are calculated is defined in a csv file. The csv file begins with the header `name,x,y,z,layer,region,type`. Each column of the file is described in Table 2. Each row corresponds to an additional electrode. The electrode csv file is read when the weights file is created, in order to populate the metadata.

**Table 2:** The format of the electrode csv file.

| Entry | Description |
| --- | --- |
| name | Either an index or a meaningful name describing the electrode |
| x | x-coordinate of the electrode (of the center of the electrode for extended electrode geometries), in $\mu\text{m}$ |
| y | y-coordinate of the electrode (of the center of the electrode for extended electrode geometries), in $\mu\text{m}$ |
| z | z-coordinate of the electrode (of the center of the electrode for extended electrode geometries), in $\mu\text{m}$ |
| layer | Cytological layer in which the electrode is located, or “NA” if in a brain region without layers, or “Outside” if not within the brain. Used only to provide reference information to the user. |
| region | Brain region in which the electrode is located, or “Outside” if not within the brain |
| type | method used to calculate the weights (Reciprocity, DipoleReciprocity, PointSource, or LineSource) |

#### 2.3.3 Generation of weights files

Discretization of the neurons into electrical compartments is performed at runtime. Therefore, to extract the number of compartments per neuron, a single-timestep simulation is run, from which a compartment report is produced. For each compartment, which are uniformly spaced, the position in Cartesian coordinates is interpolated from the 3d points in the morphology file, which do not necessarily correspond to the discretization.

After interpolation of segment positions, the H5 file defined in Section 2.3.1 is created, with all coefficients initially set to 1. In a subsequent step, coefficients are computed, e.g., based on interpolated potential field calculated using Sim4Life (ZMT Zurich MedTech AG, Zurich, Switzerland; c.f. Section A), or using the line-source approximation (c.f. Section 2.1.3) or the point-source approximation (c.f. Section 2.1.4).

While the weights files are intended for runtime calculation of extracellular signals in the Neurodamus simulation control application (see Section 2.3.4), it is also conceivable that other simulators, such as LFPy or HNN, could be extended to accept BlueRecording weights files. This would allow these simulators to use the full set of extracellular signal calculation methods that BlueRecording provides, particularly the generalized reciprocity approach, which is not available in other simulators.

#### 2.3.4 Online calculation of EEG/LFP signals

The calculation of extracellular signals is a multi-step process that begins with the launch of a Neurodamus [7] simulation. If a weights file, as defined in Section 2.3.1, is present in the SONATA simulation config file, the weights and configuration are loaded, providing the necessary information for the subsequent extracellular signal calculation.

Neurodamus then iterates through the simulation node IDs and their corresponding sections, loading the factors associated with each section from the weights file.

The registered weights are used by CoreNEURON (the compute engine used by the NEURON simulation environment [8]) to compute the extraceullular signals at run-time, once the simulation is running, by at each time-step, summing the products of the segment transmembrane currents and their corresponding segment weights.

#### 2.3.5 Reports

For storing simulation reports, the SONATA format [9] is utilized, which organizes the data in an HDF5 file (Table 2.3.5). This format is specifically designed to handle various types of data related to neural simulations, providing a structured and efficient way for storing and retrieving the data.

**Table 3:**
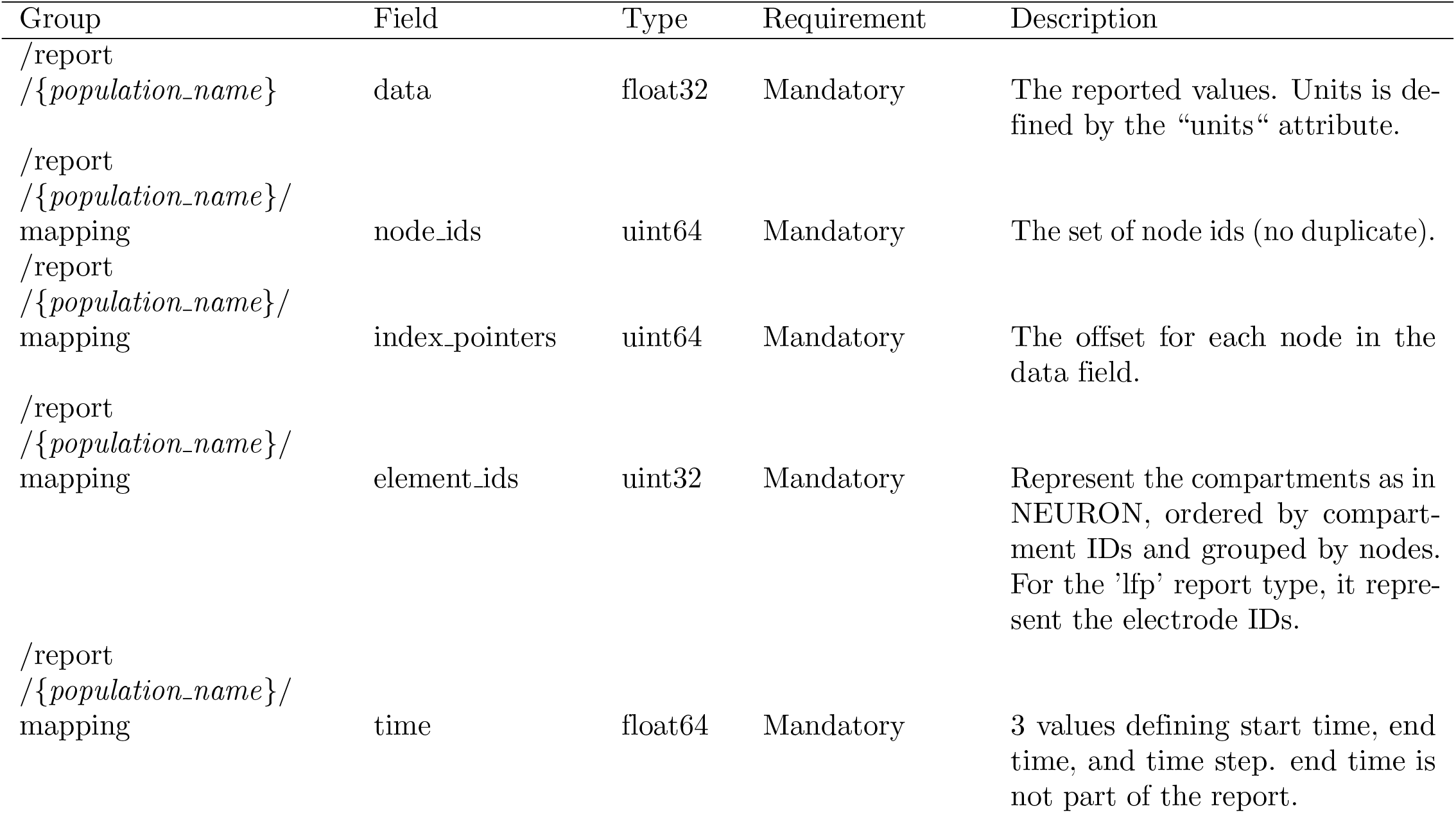
The format of the simulation reports.

**Table 4:**
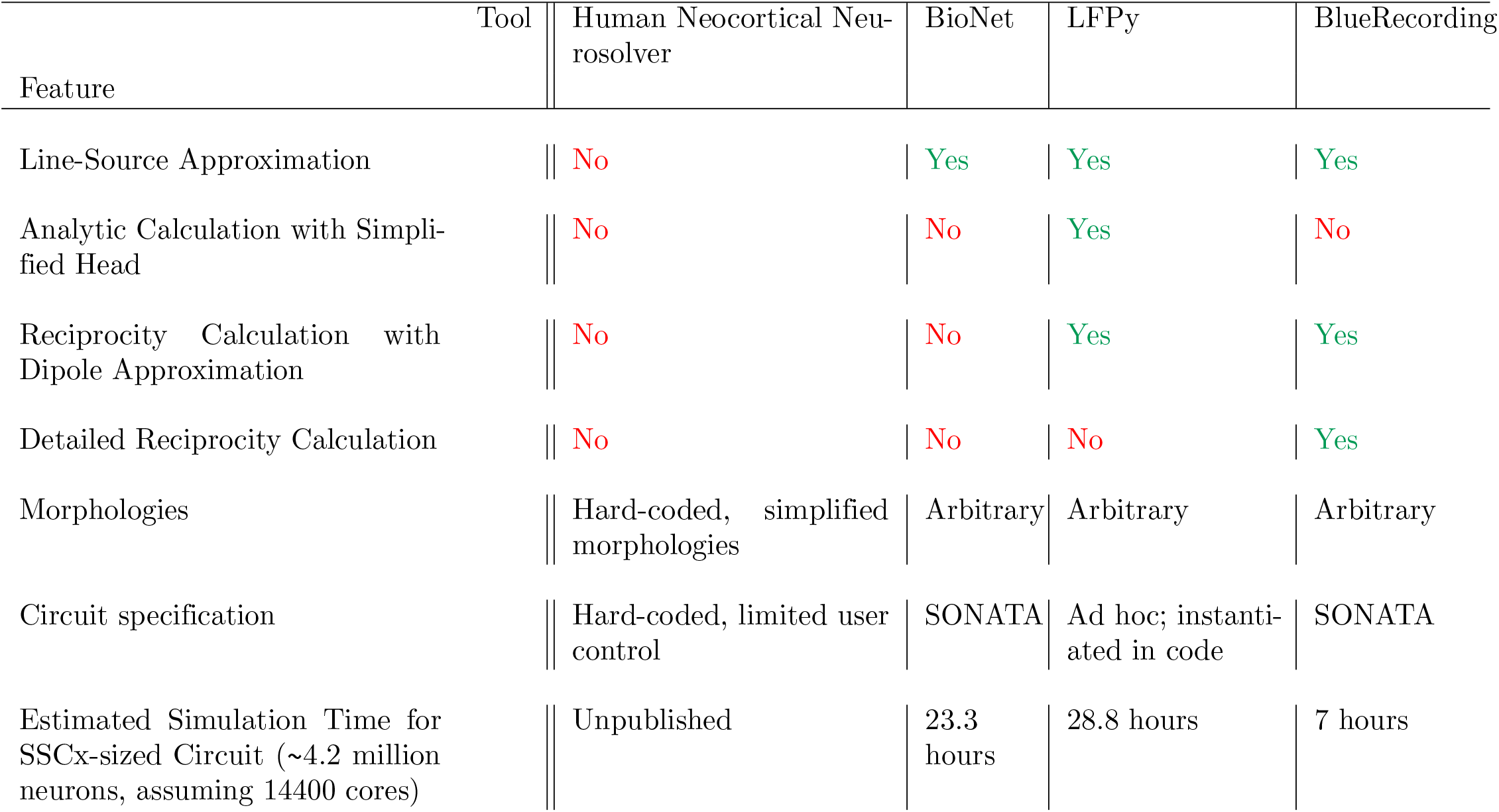
Comparison of BlueRecording with existing tools.

**Table 5:** The format of FEM output. We list only those fields which are used by BlueRecording.

| Group | Field | Size | Units | Description |
| --- | --- | --- | --- | --- |
| /Meshes/{*mesh_id*} | axis_x | $n_x$ | $m$ | x edges of mesh |
| /Meshes/{*mesh_id*} | axis_y | $n_y$ | $m$ | y edges of mesh |
| /Meshes/{*mesh_id*} | axis_z | $n_z$ | $m$ | z edges of mesh |
| /FieldGroups/{*simulation_id*}/AllFields<br>/EM E(x,y,z,f0)/_Object/Snapshots/0 | comp0 | $n_x - 1 \times n_y \times n_z$ | $V/m$ | The x-component of the E field |
| /FieldGroups/{*simulation_id*}/AllFields<br>/EM E(x,y,z,f0)/_Object/ Snapshots/0 | comp1 | $n_x \times n_y - 1 \times n_z$ | $V/m$ | The y-component of the E field |
| /FieldGroups/{*simulation_id*}/AllFields<br>/EM E(x,y,z,f0)/_Object/Snapshots/0 | comp2 | $n_x \times n_y \times n_z - 1$ | $V/m$ | The z-component of the E field |
| /FieldGroups/{*simulation_id*}/AllFields<br>/EM Potential(x,y,z,f0)<br>/_Object/Snapshots/0 | comp0 | $n_x \times n_y \times n_z$ | $V$ | The electric potential |
| / | CurrentApplied | 1 | $nA$ | Current applied between recording electrodes |

For extracellular reports, a unique aspect of the SONATA format is the use of the field “element ids” to represent electrode IDs, as opposed to representing compartments as in NEURON. This means that reports capture the electrical activity at specific electrode locations, rather than at individual compartments within the neural model.

An extracellular report is requested by adding an `lfp` block to the report section of the simulation configuration file (see [9] for a full description of this file).

~~~
“reports”: {
“report name”: {
“type”:”lfp”,
“cells”:”Mosaic”,
“variable name”:”v”,
“dt”:0.1,
“start time”: 0.0,
“end time”: 5000.0,
“sections”:”all”
}
}
~~~

The name-value pairs `“type”:”lfp”` and `“variable_name”:”v”` must be present verbatim; the others are user-configurable.

## 3 Results

### 3.1 Verification

#### 3.1.1 Unit tests

Unit tests, with 70% code coverage, are provided in our Github repository, in the folder `tests`. These ensure, for example, that the weights files are outputted with the correct format, that segment positions are interpolated correctly for a morphology with 10 sections, and that weights are calculated correctly for a simple two-compartment model. They can be run with `pytest`.

#### 3.1.2 Comparison of calculation methods

We provide a small verification example in our Github repository (https://github.com/BlueBrain/BlueRecording), comparing the generalized reciprocity approach, dipole-based reciprocity approach, line-source approximation, and point-source approximation, in a large homogeneous medium. We simulate the signal from a single Layer 5 pyramidal cell from the BBP somatosensory cortex (SSCx) model. The cell is driven with Orhenstein-Uhlenbeck noise with an amplitude of 100% of the spiking threshold and a standard deviation of 1% (c.f. Appendix B).

Based on theoretical considerations outlined in the introduction, we expect a weaker signal and disappearing differences between the approaches at larger distances. We confirm that when the recording and reference electrode are located far from the neuron (30 mm and 40 mm, respectively), the signals recorded using each of the methods are very similar (Fig. 2 A). In contrast, when the recording electrode is placed 218.5 *µ*m from the neuron, the signals recorded using the different methods begin to diverge (Fig 2 B), with the signal calculated using the dipole-reciprocity method diverging the most. This is the results of differences in the weights calculated for individual compartments (Fig 2 C, D). For the electrode closer to the neuron, the full reciprocity approach yields more positive weights (relative to the soma) for the apical tuft and more negative weights for the basal dendrites than the simplified dipole approach. The higher polarity between tuft and basals explains the larger amplitude of the signal observed.

**Figure 2:**
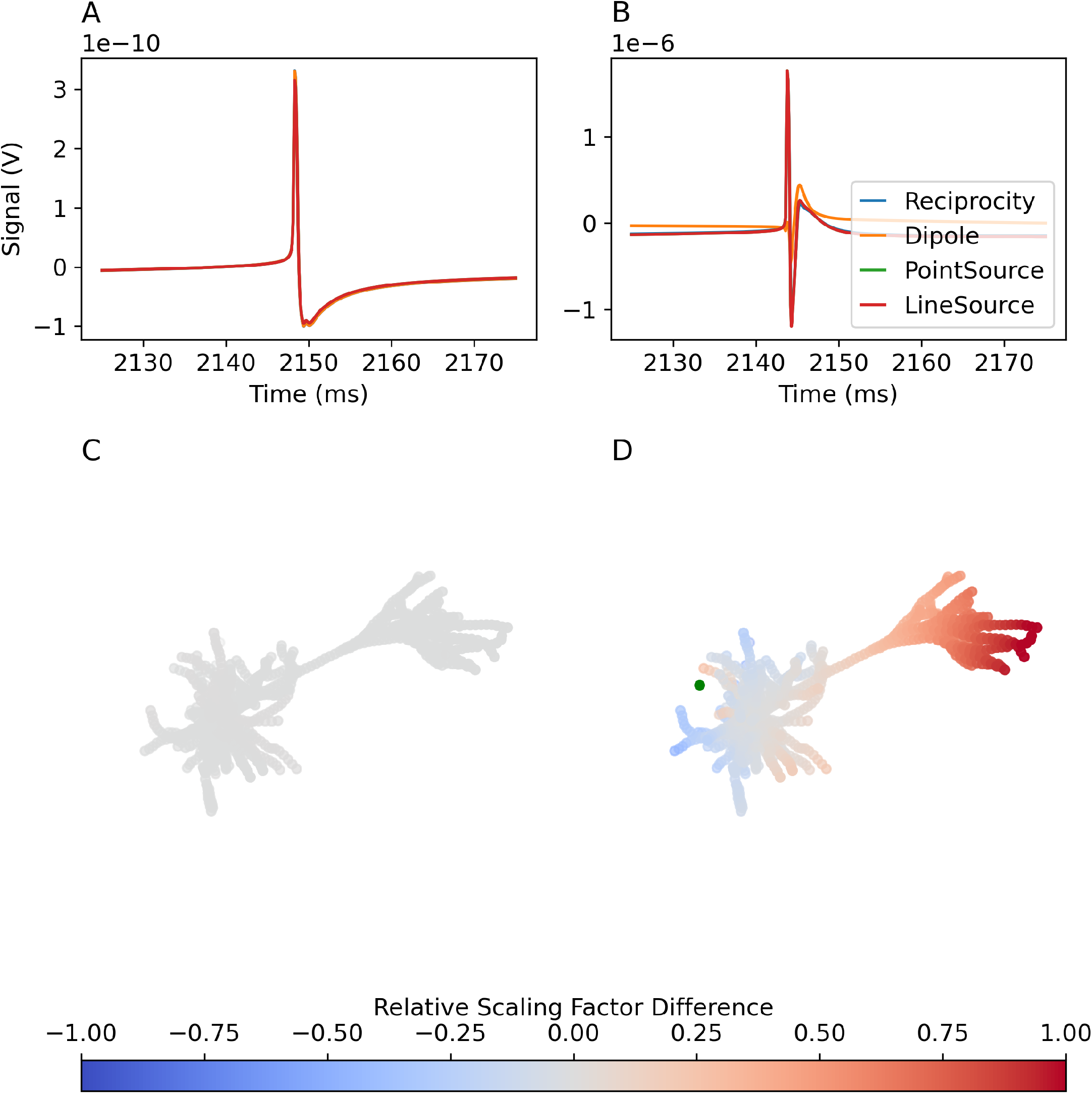
Extracellular signals recorded in a large homogeneous medium. A: Signals recorded with recording electrode (not visible in panel C) distant from the neuron. B: Signals recorded with recording electrode (green dot in panel D) near the neuron. C:Difference in per-compartment weight between generalized reciprocity and dipole-based signal calculations, for electrode far from the neuron (adjusted for constant offset, and normalized to the range of compartment weights in general reciprocity approach). D: The same, for electrode close to the neuron.

### 3.2 Biological application: Resting state EEG

We simulated a resting-state EEG, ECoG, and LFP signal originating from the Blue Brain Project reconstruction of the rat somatosensory cortex [10]. The model consists of 4.2 million biologically detailed neuron models, positioned in space according to the Paxinos Watson rat brain atlas [14], rescaled to the size of a juvenile animal. It has been shown to produce realistic firing activities [15]. In order to calculate EEG from this model, we developed a FEM model of a rat head that is spatially aligned to the BBP somatosensory cortex (c.f. Appendix A).

EEG is calculated from a recording electrode, positioned in the skull, directly above the forelimb region. A reference electrode is positioned over the hindlimb region. ECoG is produced by moving the recording electrode downward such that it is in contact with the cortical surface. (Fig 3 A). LFP is calculated by moving the electrode 1 mm into the cortex. This will place it inside cortical layer 3.

**Figure 3:**
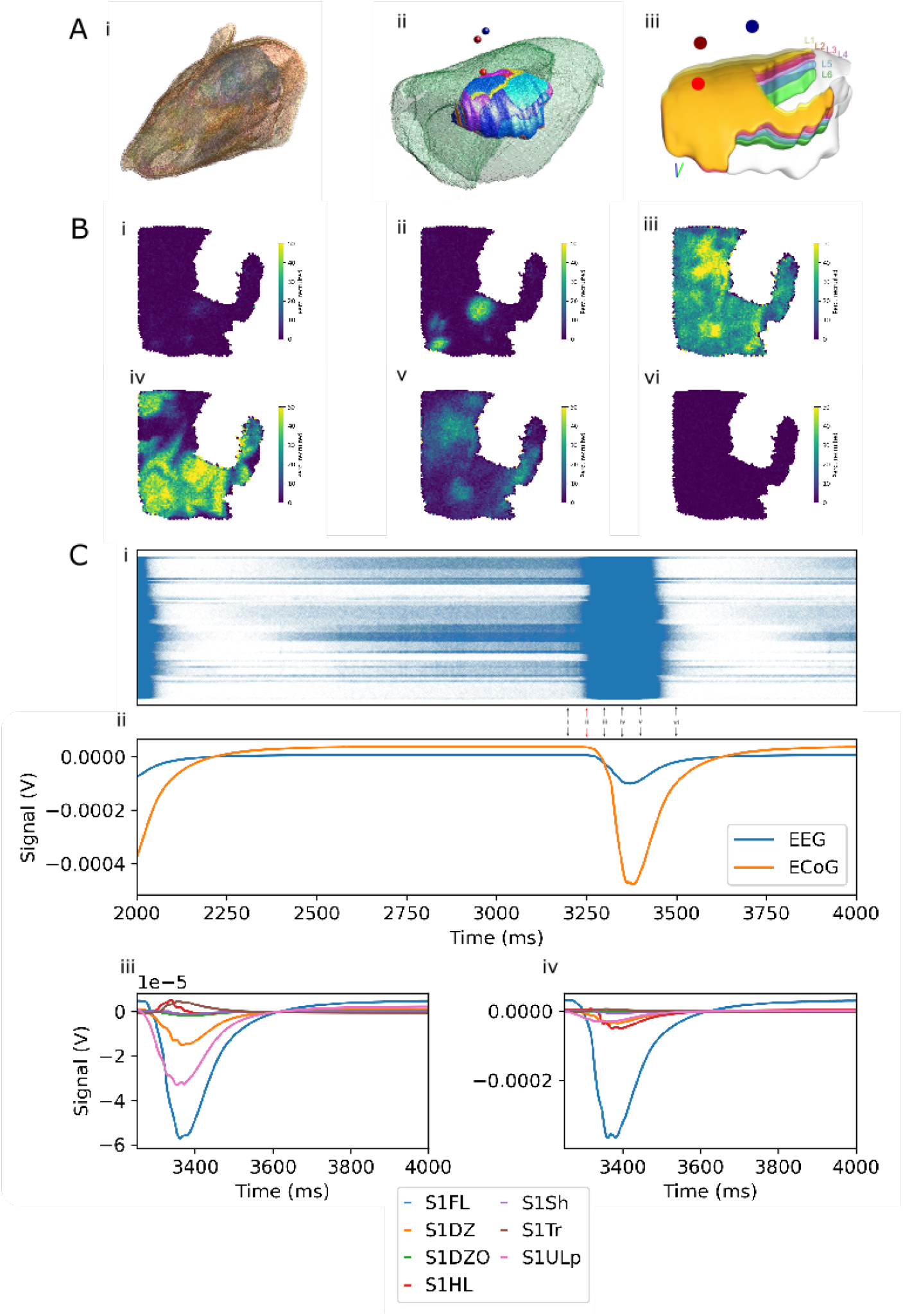
A. Rat head model. i: 3D view of the surface-mesh of the rat head model. ii: Comparison between the FEM model brain (green) and the BBP somatosensory cortex model. Locations of EEG electrodes over the forelimb region (dark red) and hindlimb region (blue) and ECoG electrode over the forelimb region (bright red) marked (not to scale). iii. Somatosensory cortex model, with approximate electrode locations as in ii. B: Top down view of mean firing rate over the somatosensory cortex at i. 3200 ms ii. 3250 ms iii. 3300 ms iv. 3350 ms v 3400 ms vi. 3500 ms. C: i: Raster plot of firing over the entire SSCx. Arrows indicate snapshots in panel B. Red arrow indicates start of window highlighted in panels C.iii and C.iv, and Figure 5. ii: EEG and ECoG recorded over the forelimb region. Green and red lines as in C.i. We note that due to baseline current noise injection, the signal is nonzero even in the absence of spiking activity. As the time course of recovery from hyperpolarization at the single-cell is longer than that of the action potential, we observe that the extracellular signal peak is broader than the firing burst. iii. Contributions of different regions of SSCx to the EEG recorded in ii. iv: Contributions of different regions of SSCx to the ECoG recorded in ii.

“Resting state” activity is achieved by injecting noisy conductance to neurons that represents uncorrelated resting-state inputs from neurons extrinsic to the model (c.f. Appendix B). The simulation is run for 5 seconds of biological time; data from the first two seconds is discarded in order to exclude an initial transient. The signal is sampled at a rate of 1 kHz. Generation of the weights file take approximately 1 hour, and the simulation runs in 7 hours on 400 nodes; each node has two 2.30 GHz, 18 core Xeon SkyLake 6140 CPUs, and 382 GB DRAM. The calculation of the extracellular signals adds no significant computational cost.

The model is simulated with an extracellular calcium concentration of 1.05 mM, resulting in bursts of activity over the entire somatosensory cortex (Fig 3B). Originating at discrete points, these bursts spread as travelling waves through the entire model (Fig 3B). They occur at a frequency of 0.66 Hz (Fig 3C.i), which is reflected in the EEG signal recorded over the forelimb region (Fig 3C.ii). The EEG signal is largely generated by the contribution of neurons from the forelimb region (FL) and the upper lip region (ULp), with smaller contributions from the dorsal zone (DZ) (Fig 3C.iii). Neurons in the hindlimb (HL) and trunk (Tr) area create a deflection with around a 10% of the amplitude, but opposite sign. The ECoG signal is still dominated by contributions from FL, with contributions from other regions significantly reduced, and without sign inversion in the HL and Tr regions (Fig 3C.iv).

To explain the difference, we investigate the weights files specifying how much the membrane current of each neuronal compartment affects the respective signal. For the EEG electrode, we found that most neurons had more positive weights associated with the apical tuft, while neurons underneath the reference electrode (corresponding to region HL) had more positive weights for their perisomatic compartments than for their tufts (Fig 4A). For ECoG, the difference in weight between apical and basal neurites is more pronounced directly under the recording electrode (corresponding to region FL), and less pronounced more laterally (corresponding to region ULp) (Fig 4B).

**Figure 4:**
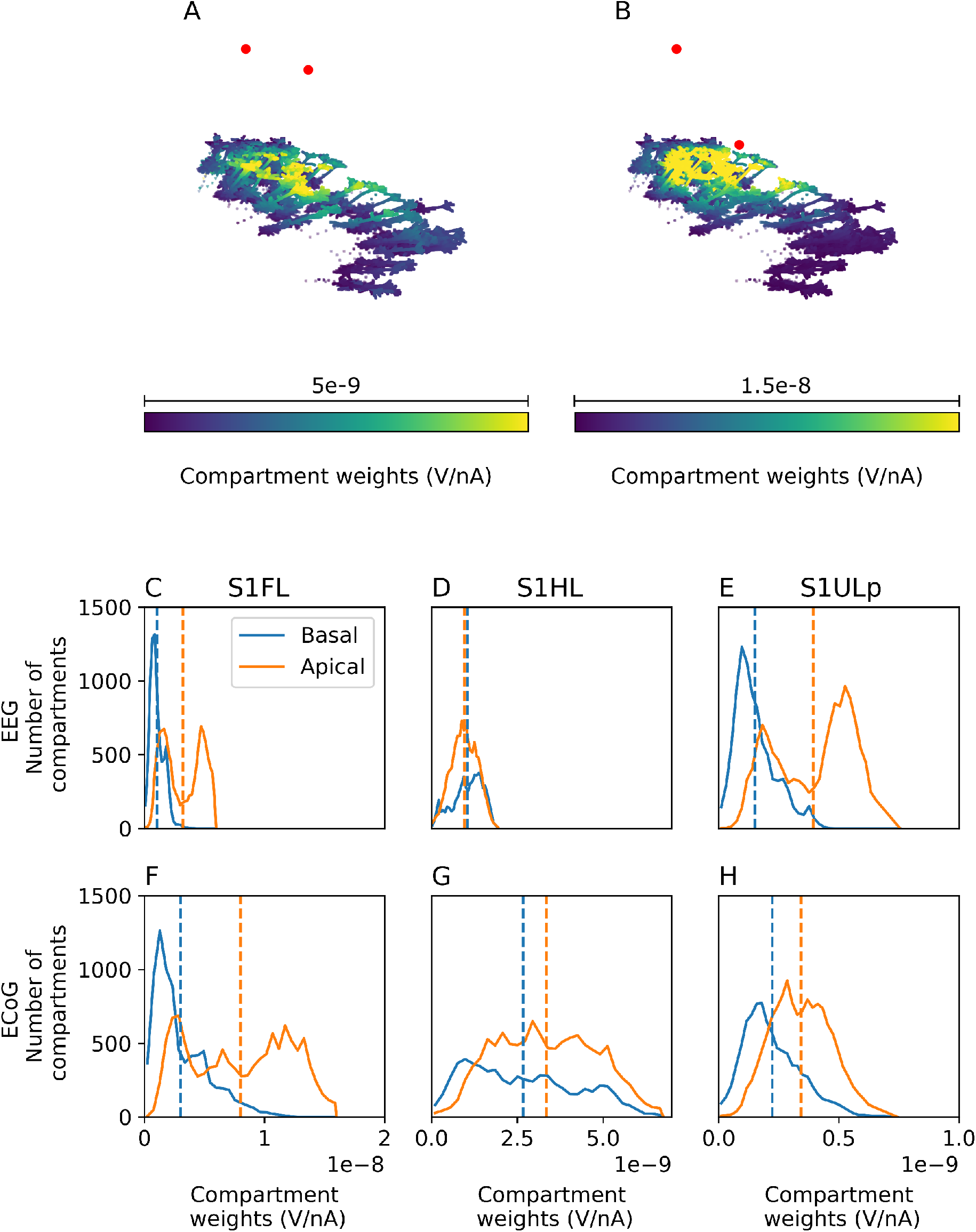
A-B: Weights for EEG and ECoG recordings, respectively, calculated using the reciprocity approach, for a sample of L5 pyramidal cells in the forelimb (compartments represented as circles), hindlimb (compartments represented as triangles) and upper lip (compartments represented as squares) regions. Electrodes are represented as red circles. Note the varying color scale ranges. C-E: Histogram of compartment weights for EEG recording, for L5 pyramidal cells in the forelimb, hindlimb, and upper lip regions, respectively. Dashed lines indicate mean values. F-H: Same as C-E, but for ECoG recording. Note that the x-axis range is different in each figure column.

We confirmed this trend by considering the distribution of weights associated with basal and apical dendrite compartments (Fig 4C-H). Specifically, apical compartments had more positive weights than basal compartments in forelimb and upper lip associated regions. For the EEG electrode, but not the ECoG electrode, basal compartments have more positive weights then apical compartments in the hindlimb region (4 D vs. G).

If membrane currents in this highly excitable state are dominated by active currents around the soma, compensated by return currents of the apical dendrites, this explains the observed inversion between contributions from the forelimb and hindlimb regions in EEG. This also demonstrates that although the EEG and ECoG signal appear very similar under the simulated conditions, their neuronal origin is very different.

#### 3.2.1 Utility of the generalized reciprocity approach

We compare the EEG, ECoG, and LFP signals obtained with the generalized reciprocity approach to those obtained with the dipole-based reciprocity approach and the line- and point-source approximations (Fig. 5 A-C). The dipole-based approach overestimates the amplitude of the EEG and ECoG signal by a factor of ∼1.5 relative to the generalized reciprocity approach (Fig. 5 A). This is attributable to differences in the contribution from the upper lip and hindlimb regions (Fig. 5 D), which can in turn be explained differences in weights due to nonlinearites in the potential field (Fig. 5 G vs J), which depend on the geometry of the dielectric environment.

**Figure 5:**
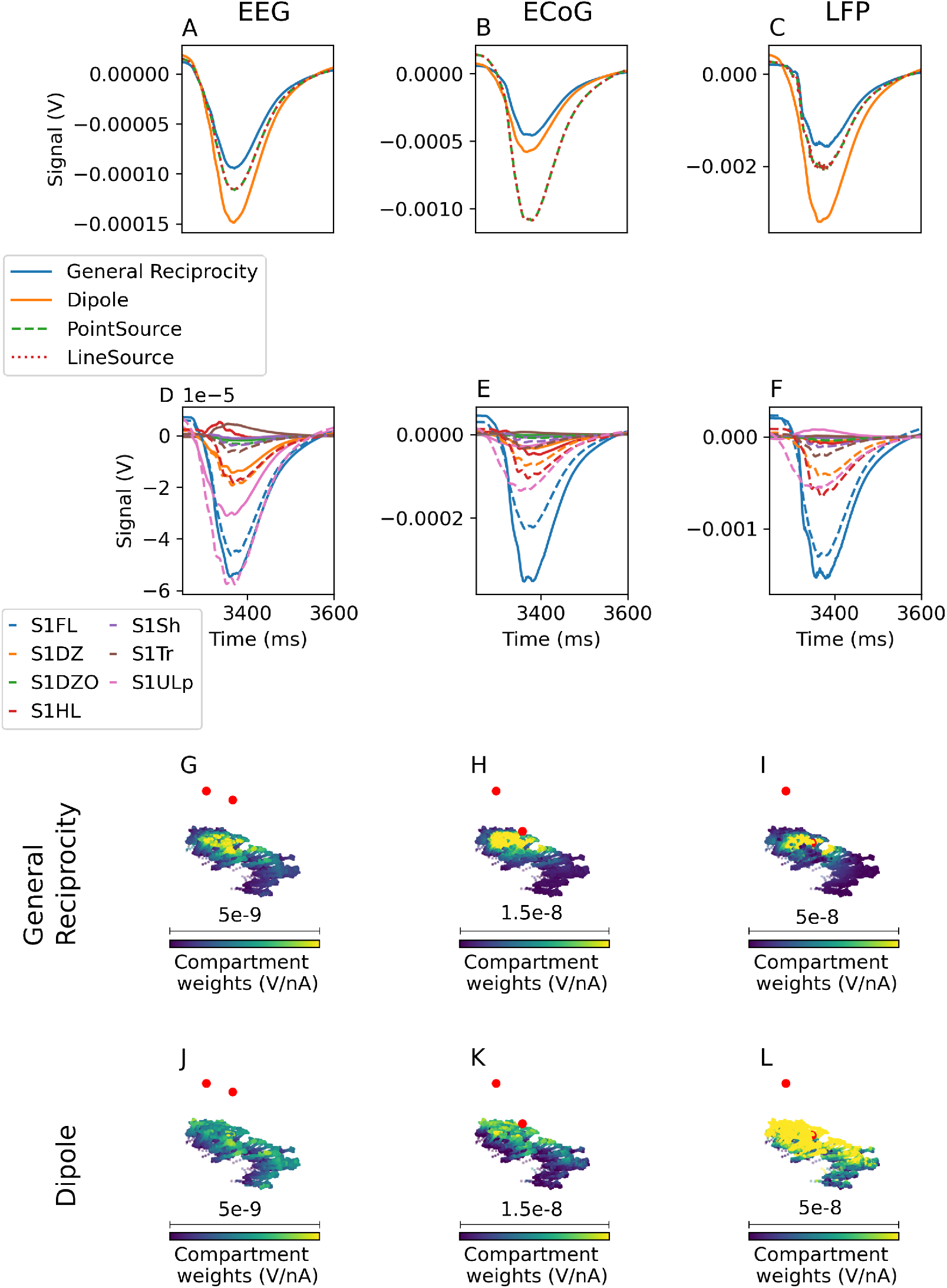
EEG (A), ECoG (B), and LFP (C) signals, recorded over, or within, the somatosensory cortex, calculated with the point-source and line-source approximation, with the generalized reciprocity theorem (ground truth), and with the simplified dipole-based approaches. D-F: Contribution of each region to EEG, ECoG, and LFP signals, respectively. Solid lines indicate general reciprocity approach, dashed lines indicate dipole approach. G-I: Weights for EEG, ECoG, and LFP recordings, respectively, calculated using the reciprocity approach, for a sample of L5 pyramidal cells in the forelimb (compartments represented as circles), hindlimb (compartments represented as triangles) and upper lip (compartments represented as squares) regions. Electrodes are represented as red circles. Note the varying color scale ranges. J-L: As in G-I, but calculated using the dipole approach

Surprisingly, the dipole approximation provides a better fit to the generalized reciprocity signal for ECoG (Fig. 5 B) than for EEG. This is because, in ECoG, the dipole approximation induces a larger error in the signal contribution from the forelimb region, in the opposite direction as the error in the contribution from the other regions (Fig. 5 E). Thus, while the general reciprocity and dipole approaches produce similar signals, their interpretation is very different.

For LFP recordings, not only does the dipole-based approach overestimate the signal amplitude – it also changes the shape of the signal, leading to a loss of information and reduced interpretability. A smaller change in shape is also observed for the point- and line-source approximations (Fig. 5 C).

While the line-source and point source approaches provide a reasonable approximation of the EEG and LFP signals obtained using the generalized reciprocity approach (Fig 5A and C), they lead to even greater error than the dipole approximation for the ECoG signal. Therefore, the generalized reciprocity approach is the only method capable of producing realistic results in all cases.

We emphasize that the dipole reciprocity approach is derived from the generalized approach under the additional simplifying assumption that the E field is constant over the range of the neural source, which is not true near the electrode (Fig. 9). The point and line source approaches similarly involve approximations that are not required by the reciprocity-based approaches, namely, that the extracellular medium is infinite and homogeneous, and that the recording electrodes are infinitesimally small. The generalized reciprocity approach, which does not rely on such simplifications, is inherently more accurate. While the line-source approach has the benefit of accounting for the finite extent of neural segments, it is evident, e.g., from Figure 3.1.2 B the the associated error is negligible, even relatively close to neurons (Fig. 3.1.2 B).

### 3.3 Biological application: Whisker flick

In a seven-column subvolume of the somatosensory cortex, we simulate a whisker flick stimulus, as in [15]. Briefly, a population of virtual (i.e., not biophysically modeled) thalamic fibers, projecting primarily to the central column of the subvolume (Fig. 6 a), is activated. averaging over 10 trials. We repeat the experiment for a circuit in which all cortico-cortical connections have been removed. EEG is recorded from the same electrodes used in the previous section.

**Figure 6:**
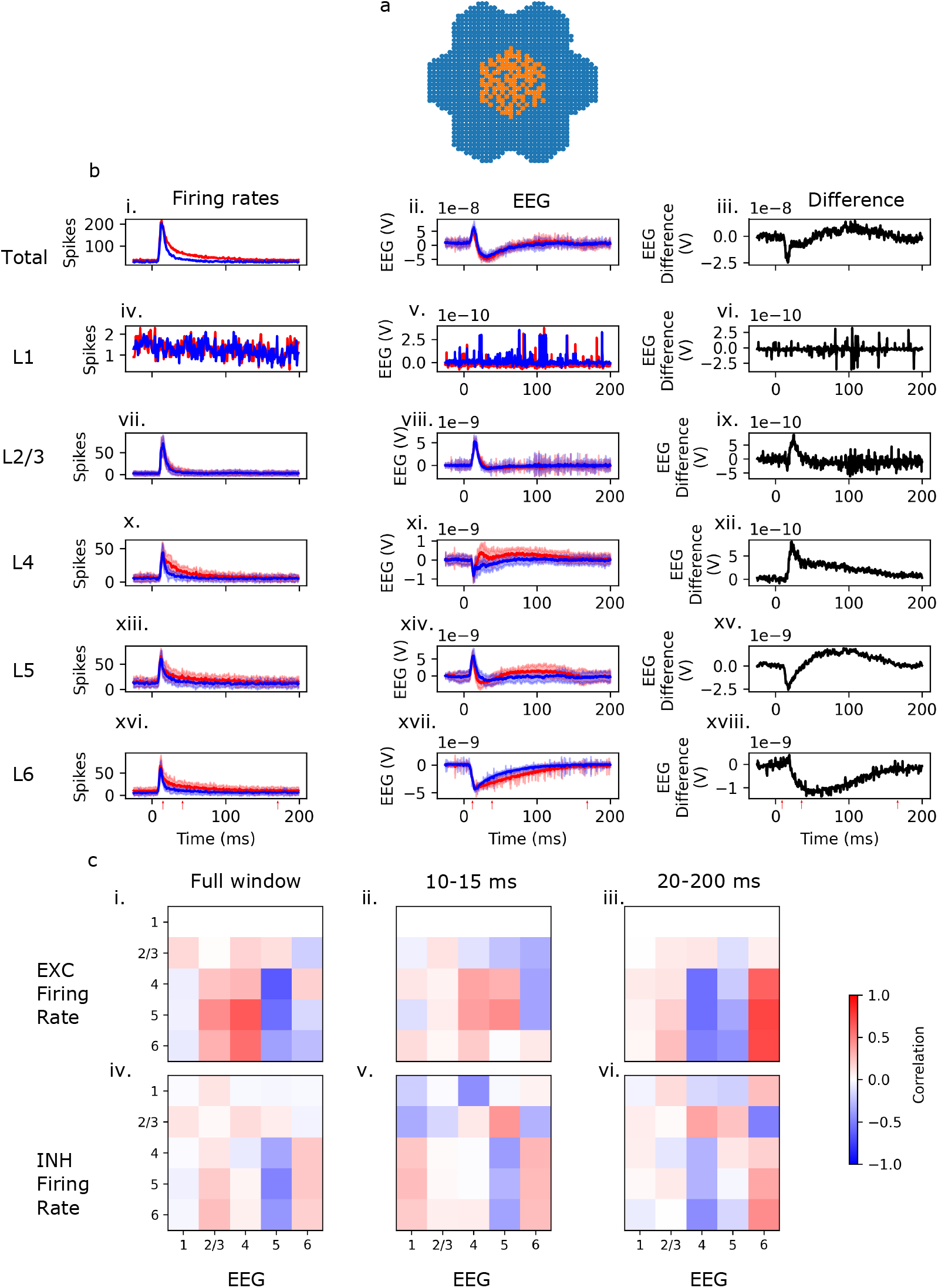
a: Selected cells from the 7-column subvolume (blue) and activated thalamic projections (orange). b: Firing rate (first column) and EEG (second column) for the original and the disconnected circuit (red and blue traces, respectively) and the difference in the EEG between the two circuits (third column), for both the full circuit (first row) and each of the layers (subsequent rows). c: Correlation matrices between excitatory (first row) and inhibitory (second row) firing rates in each layer, and the differences in EEG contributions from each layer. In each correlation matrix, firing rates are represented along the rows, and EEG differences along the columns. Correlations are calculated for the full window (first column) and for windows 12-45 ms after the stimulus, and 40-200 ms after the stimulus (second, and third columns, respectively). Start and end times of the windows are marked by red arrows in panel b.

While stimulus triggered more sustained spiking in the connected circuits than in the disconnected circuit (Fig. 6 b.i), the EEG signal is remarkably similar (Fig. 6 b.ii). Differences are observed in the EEG contributions of different layers, particularly L4 (Fig. 6 b.xi), L5 (Fig. 6 b.xiv), and L6 (Fig. 6 b.xvii). However, these differences largely compensate.

We calculate the Pearson correlation between firing rates (in the original circuit) of excitatory and inhibitory cells in each layer, and the difference between the original and the disconnected circuit in the EEG contributions of cells in each layer. Over the entire time window, excitatory firing rates tend to correlate more strongly with EEG differences than inhibitory firing rates (Fig. 6 c.i and c.v). Between 12 and 40 ms after the stimulus, inhibitory firing rates in all layers besides L2/3 correlate negatively with the difference in the L5 contribution to the EEG between the original and disconnected circuit (Fig. 6 c.v), suggesting that the negative deflection the L5 EEG difference (Fig. 6 b.xv) may be due to inhibitory inputs to L5 cells that are absent in the disconnected circuit. Between 40 and 175 ms after the stimulus, there is a positive correlation between L2/3 and L4 inhibitory firing rates and the EEG difference in Layer 5 (Fig. 6 c.vi). This suggests that the positive deflection in the L5 EEG difference (Fig. 6 b.xv) may be due to inhibitory inputs from layers 2-4 to L4 that are missing in the disconnected circuit. BlueRecording thus provides a method to suggest possible interpretations of EEG signals in terms of functional connectivity, and to predict firing rates from EEG.

### 3.4 Biological application: Hippocampal theta oscillations

As in [11], we simulated medial septal input to a subvolume of the hippocampal CA1 model consisting of a cylinder with diameter 600 *µ*m. We are also able to simulate LFP recordings from the full circuit of ∼456000 neurons. In both cases, we apply a depolarisation current of 120% of the spiking threshold to all cells. A sinusoidal current of -0.2 nA and frequency of 8 Hz is injected in PV+ cells. We mimic the effect of 1.0 uM ACh on synapses and cell excitability as described in [11]

We calculate the LFP signal in both circuits using BlueRecording, with the line-source approximation. We are able to replicate the finding in [11] that the simulated medial septal input results in LFP oscillations at ∼ 8 Hz (Fig. 7 c-g). The LFPs calculated for the two circuits (Fig. 7 d), as well as the resulting power-spectral densities (Fig. 7 e) and current-source densities (Fig. 7 f-g), are very similar in both circuits, albeit with higher power in the full hippocampal circuit (Fig. 7). These results suggest that the findings in [11] generalize to a larger circuit model.

**Figure 7:**
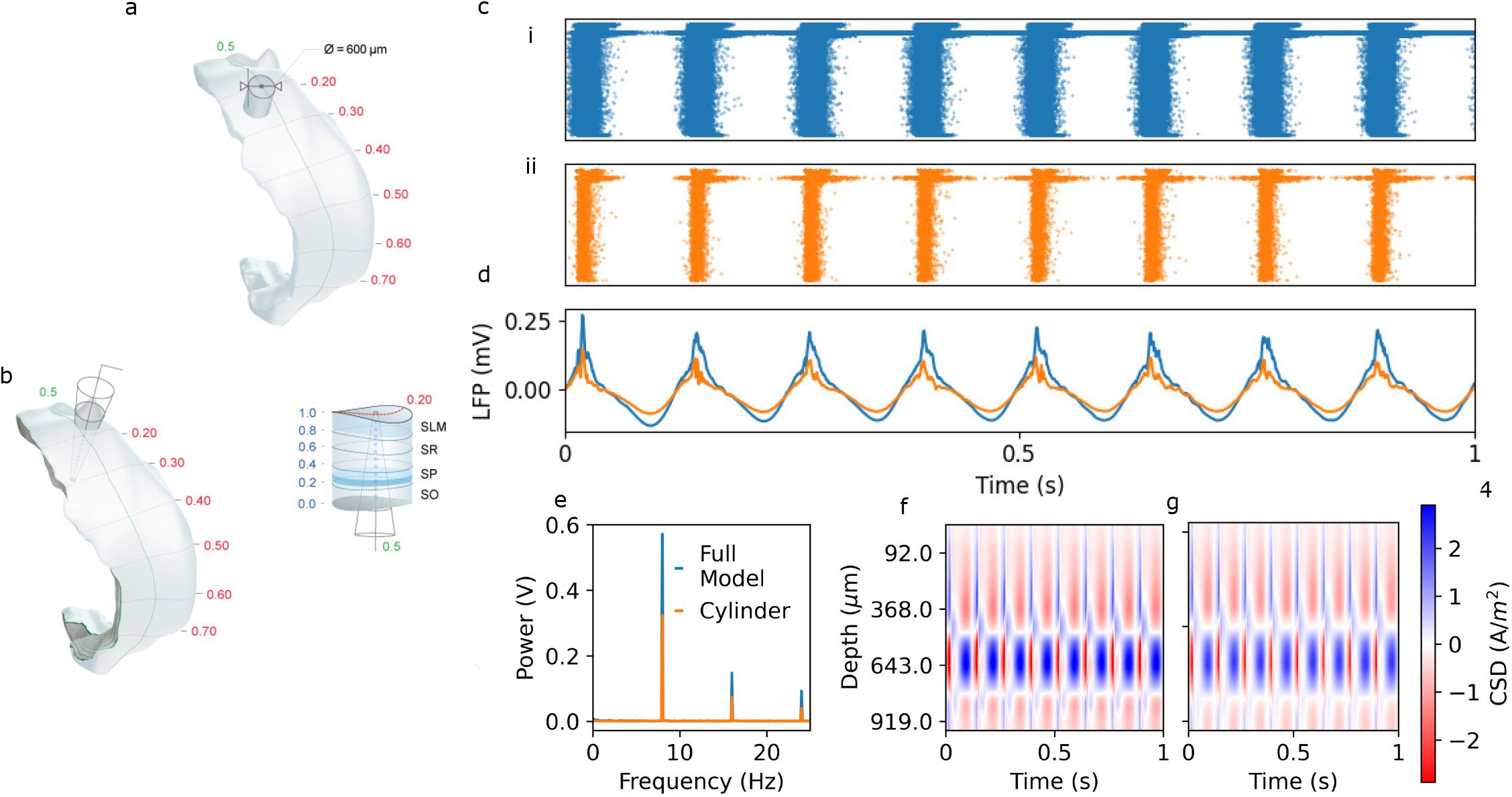
a: Visualization of the hippocampus and cylindrical subvolumes. Reproduced with permission from [11]. b: Visualization of recording electrode placement. Reproduced with permission from [11]. c: Raster plot of activity in the full hippocampus (i) and the cylindrical subvolume (ii). d: LFP recorded from a representative electrode in the hippocampus. e: Power-spectral density calculated for the signals in panel d. f: Current source density calculated in the full hippocampus simulation. g: Current source density calculated in the cylindrical circuit.

### 3.5 Comparison with existing tools

In Table 3.5, we compare BlueRecording with several existing tools. We estimate the amount of time required by these tools to run a simulation the size of the SSCx model on equivalent hardware, under the assumption that computational time is directly proportional to network size and inversely proportional to the number of available CPU cores. Thus, for a network of size *N* cells run on *C* cores, computational time 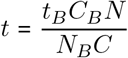, where *t*_*B*_, *N*_*B*_, and *C*_*B*_ are reported times, network sizes, and numbers of cores reported in [16] and [17] for LFPy and Bionet, respectively. We do not account for differences in CPU clock speed or available RAM. We show that both in terms of available signal calculation methods and in terms of performance, BlueRecording outperforms the state of the art by a significant margin. However, we were unable to assess the performance of HNN, and we note that LFPy supports several analytical methods for the calculation of extracellular signals that are not supported by BlueRecording. Because the calculation of extracellular signals itself is not a significant source of computational cost, either in BlueRecording or in the alternatives described above, the performance advantages of BlueRecording can be attributed to its tight integration with CoreNEURON, which is optimized for computational performance for large networks. In contrast, LFPy depends on a Python API that induces significant overhead.

## 4 Availability and Future Directions

### 4.1 Code availability and dependencies

- The source code for BlueRecording, and instructions for generating the figures in Sections 3.1.2 and 3.2 are available at https://github.com/BlueBrain/BlueRecording.
- The seven column subvolume of the BBP circuit model is available under the following DOI: 10.5281/zenodo.7930276. The full BBP circuit model will be made available upon request.
- Membrane mechanism files used in the neuro-simulations are available at https://github.com/BlueBrain/neurodamus-models.
- The specification for the version of the SONATA format used at BBP is available at https://github.com/BlueBrain/sonata-extension/.
- Neurodamus is available at https://github.com/BlueBrain/neurodamus.
- CoreNEURON itself is fully integrated into the NEURON simulation environment, which is available at github.com/neuronsimulator/nrn.
- Code for the generation of FEM models, as well as a list of dependencies, is available at https://github.com/BlueBrain/BlueBrainHeadModels. The required finite element meshes are available on Zeonodo (DOI: 10.5281/zenodo.10926947). As this pipeline is specific to the demonstration application presented in this work, we do not consider it to be part of the core BlueRecording package.
- Finite element meshes and FEM output files used in the simulations in Sections 3.1.2 and 3.2 are available on Zenodo (DOI: 10.5281/zenodo.10927050)

### 4.2 Long-term outlook

We anticipate that BlueRecording can be easily extended to permit simulation of linear signals from other recording modalities, both electromagnetic, such as MEG, and optical, such as VSDI, by simply writing the appropriate coefficients to the weights file, and in the case of VSDI, multiplying the weights by the segment voltage rather than the transmembrane current. A weights file could also be generated to directly calculate the current source density, or the current dipole itself, rather than an electromagnetic signal.

Because BlueRecording is compatible with the SONATA format, it can readily be used in hybrid models, where the dominant contributors are modeled in morphological detail, while other contributors are represented by simplified point neurons (similar to HNN) or neural masses. Such hybrid models permit the simulation of even larger models, or to run SSCx-sized models on less performant computational infrastructure.

## Acknowledgements

We thank Julian Budd for his assistance with the hippocampus model, and Stephanie Jones for her helpful comments on the paper. This work was supported by funding to the Blue Brain Project, a research center of the École polytechnique fédérale de Lausanne (EPFL), from the Swiss government’s ETH Board of the Swiss Federal Institutes of Technology.

## A Supplementary methods: Finite-element models

In order to perform the FEM electromagnetic simulations required for the reciprocity-based approach, we must generate a FEM model of the rat head that is spatially aligned to the BBP microcircuit, which is based on the Paxinos Watson atlas. We begin the process of generating the FEM model with the series of segmented MRI images underlying the ViZOO NeuroRat (150g) model V4.0 (DOI: 10.13099/VIP91106-04-1, https://itis.swiss/virtual-population/animal-models/animals/neurorat/). The procedure used for aligning the NeuroRat model to the BBP circuit model is outlined in Fig. 8.

**Figure 8:**
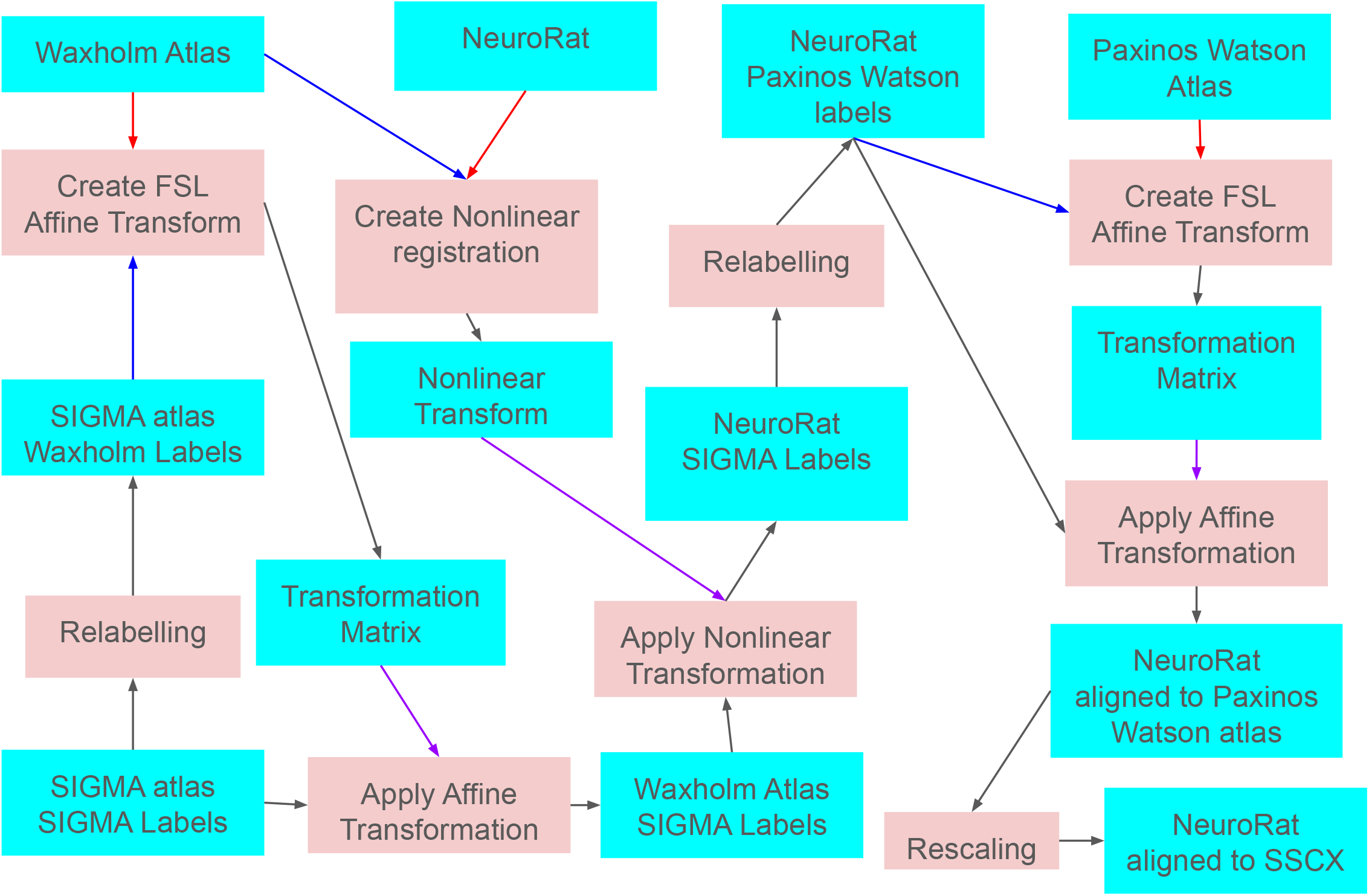
Procedure for aligning the NeuroRat head model underlying the FEM simulations with the BBP circuit model. Blue blocks represent input and output files, while pink blocks represent processes. For processes that generate transformations between two images, blue arrows represent the moving image, while red arrows represent the target image. Black arrows represent inputs that are transformed by processes and the resulting outputs. For processes that apply a transformation to an image, purple arrows represent the transformation object used.

**Figure 9:**
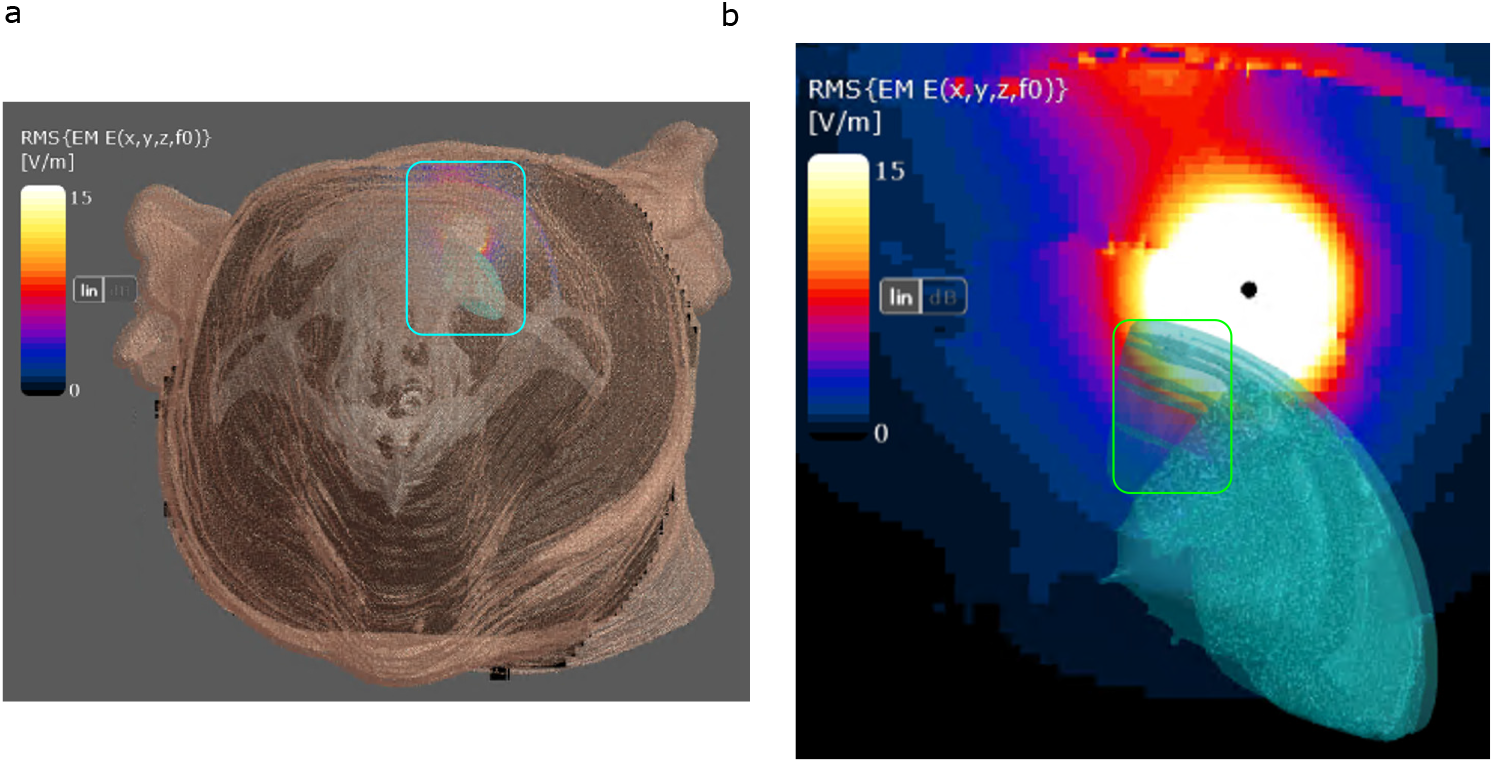
a. E-Field magnitude, in a cross section of the rat head, induced by current applied to an ECoG recording electrode. The scalp (beige), cancellous skull (white), and somatosensory cortex (blue) are displayed for context. b. Zoom in on the boxed area in panel a, with scalp and skull removed. Somatosensity cortex is displayed in blue. The E-field magnitude within the somatosensory cortex varies significantly (highlighted in the green box), demonstrating that the dipole approximation is not valid for ECoG recordings.

The NeuroRat head model is segmented according to the Waxholm atlas [18]. As this brain segmentation is relatively coarse, we re-segment the NeuroRat brain with labels from the SIGMA atlas [19]. In order to do so, we create a version of the SIGMA label map, with each region assigned to the corresponding coarser labels from the Waxholm atlas. An affine transformation between the relabelled SIGMA label field and the Waxholm atlas is calculated using the FSL FLIRT tool with a label-difference metric [20]. This transform is applied to the original SIGMA model, yielding a model that is aligned to the Waxholm atlas but with SIGMA labels.

A nonlinear transformation aligning the Waxholm atlas to the NeuroRat is calculated with Advanced Normalization Tools (ANTs). This transforamtion is applied to the Waxholm atlas but with SIGMA labels, yielding a model aligned to the NeuroRat, but with SIGMA labels. The SIGMA labels are assigned to the corresponding labels in the Paxinos Watson atlas, and an affine transform is calculated, as before, from the NeuroRat brain to a digitization of the Paxinos Watson atlas created, as described in Supplementary Materials A of [21], by aligning, rasterizing and interpolating the individual slices obtained from the CD-ROM distributed with [14]. This transformation is then applied to the full NeuroRat label field (cropped to include just the head), to produce a rat head model aligned to the Paxinos Watson atlas. A scaling factor of 0.96837 is then applied to account for the difference between the adult and juvenile rat.

Finally, the label field is transformed into a discretized mesh (Fig 3 A i.) using Sim4Life (ZMT Zurich MedTech AG). The mesh positioning is manually adjusted to ensure that the upper layers of the SSCx does not extend into the cerebrospinal fluid. This procedure resulted in a good match between the Paxinos Watson brain and the FEM head model (Fig. 3 A. ii.). Recording electrodes, modelled as small spheres, are positioned on the head model as described in Section 3.2.

FEM simulations were executed using the Electro-Ohmic Quasi-Static solver in Sim4Life, which solves the equation ∇*σ*∇*ϕ* = 0, where *σ* is the electrical conductivity and *ϕ* is the electric potential, from which the electric field can be obtained as *E* = −∇*ϕ*. This quasi-static approximation of Maxwell’s equations can be used, because at the frequencies of interest, displacement currents are negligible compared to ohmic currents and the domain is much smaller than the wavelength. Tissue properties were assigned in accordance with the low-frequency dielectric properties from the IT’IS Tissue Properties Database V4.1 [22]. All brain regions outside of the cerebellum and brainstem are assigned a conductivity of ∼0.37S/m, and both cortical and cancellous skull are assigned a conductivity of ∼0.018S/m. To determine the electric potentials required for the application of the general form of the reciprocity theorem, Dirichlet boundary conditions were applied to one of the recording electrodes (1 V) and the reference electrode (−1 V). The applied total current is calculated by integrating the normal component of the current flux density *j* = *σE* over a closed surface surrounding the recording electrode, but excluding the reference electrode. The cortex is discretized at a resolution of 0.2 mm, the skull at a resolution of 0.4 mm; the electrodes are discretized using the Sim4Life Automatic Grid at Extremeley Fine resolution, while other tisses are discretized at Default resolution. This results in a mesh of ∼30 MCells, with a minimum resolution of ∼20*µ*m and a maximum resolution of ∼1cm. The solver convergence settings are set to a relative tolerance of 1e-12, an absolute tolerance of 1e-50, a divergence value of 1e50, and a iteration number maximum of 100000.

## B Supplementary Methods: Noise input

In [15], noisy conductance was injected into the neurons of the SSCx model in order to represent uncorrelated synaptic input. This was accomplished in Neurodamus using the `SEClamp` mechanism built into neuron, which models a voltage clamp with variable input resistance. For numerical reasons, the currents injected by the `SEClamp` are calculated at different time points than the transmembrane currents, meaning that total currents do not obey Kirchoff’s law, leading to inaccurate calculations of extracellular signals. We therefore created a new mechanism, `conductanceSource`, which is identical to `SEClamp`, but which is treated by CORENEURON as an ion channel current rather than an electrode current. By using `conductanceSource` instead of `SEClamp` to inject noise, the input current can be accounted for in the calculation of the extracellular signals. Similarly, we add a `MembraneCurrentSource` mechanism to replace the Neuron builtin `IClamp` current clamp.

As there may be cases in which a user wishes to use `SEClamp` or `IClamp` to model a physical voltage clamp rather than a synaptic input, we add the key `represents physical electrode` to the SONATA simulation configuration file for all noise sources. If this key is set to True, the `SEClamp` or `IClamp` mechanism is used, for conductance and current sources, respectively. In these cases the injected current will not be accounted for in the calculation of the extracellular signal. For example, the relevant block of the SONATA simulation configuration file for a physical electrode injecting noisy conductance might read as follows

~~~
“Stimulus gExc L23E”:{
“input type”: “conductance”
“module”: “relative ornstein uhlenbeck”,
“delay”: 0,
“duration”: 3000,
“reversal”: 0,
“tau”: 2.7,
“mean percent”: 17.918,
“sd percent”: 7.167,
“node set”: “Layer23Excitatory”,
“represents physical electrode”:true
}
~~~

## C Supplementary Methods: FEM output file format

BlueRecording assumes that FEM output files used to calculate compartment weights are formatted in the same way as output files from a Sim4Life Electro-ohmic Quasi-static simulation. The format of the file is described in Table C:

## D Supplementary results: E field variability in the cortex

